# Transfer Learning for Predicting Conversion from Mild Cognitive Impairment to Dementia of Alzheimer’s Type based on 3D-Convolutional Neural Network

**DOI:** 10.1101/2019.12.20.884932

**Authors:** Jinhyeong Bae, Jane Stocks, Ashley Heywood, Youngmoon Jung, Lisanne Jenkins, Aggelos Katsaggelos, Karteek Popuri, M. Faisal Beg, Lei Wang, for the Alzheimer’s Disease Neuroimaging Initiative

## Abstract

Dementia of Alzheimer’s Type (DAT) is associated with a devastating and irreversible cognitive decline. As a pharmacological intervention has not yet been developed to reverse disease progression, preventive medicine will play a crucial role for patient care and treatment planning. However, predicting which patients will progress to DAT is difficult as patients with Mild Cognitive Impairment (MCI) could either convert to DAT (MCI-C) or not (MCI-NC). In this paper, we develop a deep learning model to address the heterogeneous nature of DAT development. Structural magnetic resonance imaging was utilized as a single biomarker, and a three-dimensional convolutional neural network (3D-CNN) was developed. The 3D-CNN was trained using transfer learning from the classification of Normal Control and DAT scans at the source task. This was applied to the target task of classifying MCI-C and MCI-NC scans. The model results in 82.4% classification accuracy, which outperforms current models in the field. Furthermore, by implementing an occlusion map approach, we visualize key brain regions that significantly contribute to the prediction of MCI-C and MCI-NC. Results show the hippocampus, amygdala, cerebellum, and pons regions as significant to prediction, which are consistent with current understanding of disease. Finally, the model’s prediction value is significantly correlated with rates of change in clinical assessment scores, indicating the model is able to predict an individual patient’s future cognitive decline. This information, in conjunction with the identified anatomical features, will aid in building a personalized therapeutic strategy for individuals with MCI. This model could also be useful for selection of participants for clinical trials.

## I. INTRODUCRION

**D**EMENTIA of Alzheimer’s Type (DAT) is a common and severe neurodegenerative disorder [2, 14]. Current research in pharmacological intervention for DAT has not yet been able to reverse the disease course. Therefore, Mild cognitive impairment (MCI), as a precursor to dementia, is a crucial area for research as a potential point of intervention. MCI patients are characterized by noticeable cognitive decline, including deficits in memory or language. Critically, a patient with MCI can progress into DAT or remain stable in their MCI diagnosis. About 10%∼12% of MCI patients convert to DAT every year [31].

Predicting patients who progress from MCI to DAT is important for patient care, as well in the selection for clinical trials aimed at treating and preventing disease [32]. However, diagnostic tools for DAT, which rely heavily on clinical scores, are limited in their ability to predict future development of the disease. Thus, new methodology is needed in order to better predict disease progression.

With the development of computational methods such as machine learning and deep learning, the utility of biomarker-based diagnosis for the classification and prediction of disease is becoming recognized. Various methods have been proposed to tackle the problem of predicting MCI patients who convert to DAT (MCI-Converters or MCI-C) vs. those who do not (MCI-Non-Converters or MCI-NC) [4, 7, 21, 38]. For example, using Random Forest with weak hierarchical lasso feature selection, Li et al. [21] achieved 74.8% classification accuracy with 161 MCI-NC and 132 MCI-C sMRI scans. Cheng et al. [7] produced 79.4% classification accuracy by using Domain Transfer Feature Selection (DTFS) and Domain Transfer Sample Selection (DTSS) for extracting features and Support Vector Machine (SVM) for classifying 43 MCI-NC and 56 MCI-C patients. Similarly, Suk et al. [38] had 74.8% classification accuracy in classifying 226 MCI-NC and 167 MCI-C patients by using 2D-CNN based on 93 regions of interest (ROIs) as features.^1^ Lastly, Basaia and colleagues [4] showed 74.9% classification accuracy in classifying 533 MCI-NC and 280 MCI-C patients by using 3D-Convoluational Neural Network (CNN) based on gray matter tissue probability maps.

There are several limitations to the previous studies described so far. For example, many of these failed to assess their model using a separate, independent test dataset, which is the best practice in the field to evaluate a model’s effectiveness and generalizability [33], particularly in the absence of feature visualization. When designing a study, it is important to assign a portion of the whole data set in a random manner to be included in the independent test set [19]. For example, Basaia and colleagues [4] assigned 10% of the whole dataset as test. While the most effective splitting ratio of the training, validation, and test sets is still under discussion, the ratio of 60:20:20 or 70:15:15 is traditionally accepted for small data set.

In addition, previous methods relied on hand-crafted feature extraction, whereby raw data are transformed to produce specific features (e.g., cortical thickness) that train the machine [1]. This approach assumes that the chosen feature is the most informative but may miss important information contained in the raw data. For example, studies that selected gray matter as the feature for model training [4] did not consider CSF or white matter that also play a role in DAT [17, 20, 40]. Additionally, Cheng et al. [7] manually selected “useful” samples using DTFS and DTSS-based features. Machined trained with such samples may be biased and thus may not be generalizable to other populations.

CNN is a deep-learning approach that has evolved in recent years to produce better classification performance than conventional machine learning methods across several fields [6]. An end-to-end CNN is also able to produce features that are not biased to the researcher’s choice. However, this has not been implemented to predict conversion from MCI to DAT.

## II. RESEARCH OBJECTIVES

In the present study, we implement an end-to-end CNN model with transfer learning [29] to classify MCI-NC vs. MCI-C patients using structural magnetic reasoning image (sMRI). We evaluate model performance up to 10 years before conversion. Further, using an occlusion map method for visualization, we determine which regions of the brain are most significant in the prediction model. Finally, we correlate model prediction probability with diagnostic and clinical measures.

## III. BACKGROUND INFORMATION

Transfer learning improves model performance by training the model through two classification tasks, i.e., the source task and the target task. At the source task, the model is pre-trained with the resource that is similar to the target task. Through the source task, domain knowledge is generated, and it is transferred to the target task. The model is re-trained with the resource that is directly relevant to the classification objective based on the domain knowledge. This scheme enables the model to be optimized more efficiently.

Previous research suggests that the classification task of Cognitively Normal Control (NC) vs. DAT is similar to the classification task of MCI-NC vs. MCI-C [8, 9, 43]. In previous studies, the classification task of NC vs. DAT has been used to pre-train the model [4, 7]. Therefore, we utilize a classification task of NC vs. DAT as the source task for transfer learning to our target task model.

Visualizing features that are significant in the model’s predictions is important as it enables us to validate the model’s reasoning. It also allows for identification of neuroimaging biomarkers of conversion to DAT. State-of-the-art visualization techniques include Gradient Class Activation Map (Grad-CAM) and Guided Gradient CAM (Guided-Grad-CAM) due to its class-discriminative nature of feature visualization [34, 45]. However, in the medical field, this could act as a drawback as it could not visualize the characteristic of negative samples [3]. Also, the visualized features have extremely low resolutions (e.g., 3×4×3 voxels) due to the CNN architecture design. In occlusion method [44], small sections (e.g., 2×2×2 voxels) are systematically occluded from all scans, and the already-trained model produces prediction scores on the occluded scans, producing an occlusion map which represents the prediction scores for the occluded image location. The resulting map of relevant brain regions is at a resolution that is close to the original data. To the best of our knowledge, this method has yet to be implemented in the deep model of classifying MCI-NC vs. MCI-C.

## IV. MATERIALS AND METHODS

### A. Participants

Data used in the preparation of this article were obtained from the Alzheimer’s Disease Neuroimaging Initiative (ADNI) database (adni.loni.usc.edu). The ADNI was launched in 2003 as a public-private partnership, led by Principal Investigator Michael W. Weiner, MD. The primary goal of ADNI has been to test whether serial magnetic resonance imaging (MRI), positron emission tomography (PET), other biological markers, and clinical and neuropsychological assessment can be combined to measure the progression of mild cognitive impairment (MCI) and early Alzheimer’s disease (AD).

The source task includes 2084 NC and 1406 DAT scans from 1080 subjects. Scans from multiple timepoints are included if available. In the target task, we examine MCI-C patients with a conversion time up to 3 years (longer conversions are examined later), and MCI-NC patients with a duration of MCI (within ADNI) at least 3 years. MCI subjects with a duration of MCI less than 3 years without conversion are exclude due to the potential possibility of near-future conversion. Only single timepoints are included for the target task, resulting in 222 MCI-NC and 228 MCI-C scans from 450 subjects. Demographic information and clinical scores are shown in Table 1. Figure 1 shows the distributions of duration of MCI-NC and conversion time of MCI-C patients.

**Table 1.** Demographic and clinical information within subjects for the Source and Target Tasks.

|  |  | N<sub>total</sub> | Age | %<br>Male | Education | CDRSB | ADAS11 | ADAS13 | MMSE |
| --- | --- | --- | --- | --- | --- | --- | --- | --- | --- |
| *Source Task* | NC | 2084 | 76.49<br>(5.92) | 49.8% | 16.35<br>(2.74) | 0.09<br>(0.30) | 5.56<br>(2.85) | 8.70<br>(1.32) | 29.04<br>(1.21) |
|  | DAT | 1406 | 76.18<br>(7.22) | 60.1% | 15.35<br>(2.90) | 5.22<br>(2.41) | 20.47<br>(7.85) | 31.03<br>(9.43) | 22.31<br>(3.68) |
| *Target Task* | MCI-NC | 222 | 72.25<br>(7.32) | 63.1% | 15.97<br>(2.85) | 1.18<br>(0.63) | 8.61<br>(3.41) | 13.77<br>(5.33) | 28.00<br>(1.69) |
|  | MCI-C | 228 | 74.18<br>(6.96) | 57.0% | 15.87<br>(2.78) | 1.97<br>(0.98) | 13.17<br>(5.00) | 21.27<br>(6.04) | 26.77<br>(1.72) |

Results are reported as mean (SD). Age and education are reported in years. CDR=Clinical Dementia Rating Scale; ADAS11=Alzheimer’s Disease Assessment Scale 11; ADAS13=Alzheimer’s Disease Assessment Scale 13; MMSE=Mini Mental State Exam.

**Figure 1.**
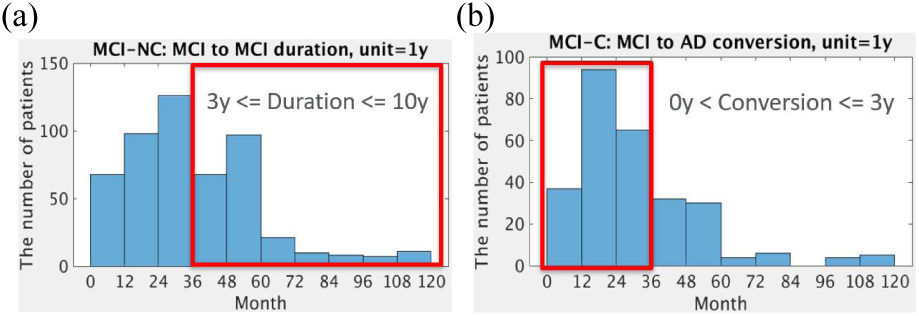
Distribution of MCI-NC (N=514) and MCI-C (N=277) patients according to the duration and conversion years. As the classification task lies in classifying MCI in patients who would convert to DAT in 3 years, MCI-C patients whose conversion time is in 3 years (red box in (b), N=228) are included. As a comparison to this group, MCI-NC patients whose duration time is at least 3 years (red box in (a), N=222) are included in this study.

### B. Structural MRI data preprocessing

1.5T and 3T sMRI data are downloaded from ADNI. Preprocessing is performed including skull-stripping [41], re-orientation, cropping and padding to 158 × 196 × 170. The FMRIB Software Library (FSL; https://fsl.fmrib.ox.ac.uk) is then used to correct intensity inhomogeneity by using an N3 algorithm [37] and to co-register the scans to the Montreal Neurological Institute (MNI)-152 atlas by using affine linear alignment.

### C. Data setup for transfer learning

For the source task, NC and DAT scans are randomly selected and divided into training, validation, and test sets. To provide diverse domain knowledge as much as possible, 90% of the data (3143 scans) are assigned to the training set while the validation and test sets each contained 5% of the sample (172 and 175 scans). Groups within the training, validation and tests sets are confirmed not to differ significantly on demographic and clinical characteristics: sex, race, ethnicity, marital status, age, years of education, clinical scores, genetic information, etc ^2^.

For the target task, MCI-NC and MCI-C scans are split into training, validation, and test set by following the conventional ratio of 70% vs. 15% vs. 15% (314, 68, and 68 scans). To avoid data leakage [42], a single time point scan is used for each subject. The test set of the target task is also ensured to be fully independent (i.e., unseen) from the training and validation sets in both the source and target tasks. Therefore, no subjects in the target task test set overlap with the rest of the sample. This step has been overlooked in previous research and is crucial for both avoiding biased learning and increasing the generalizability of the model.

### D. Architecture of Convolutional Neural Network

A base model for transfer learning is developed by benchmarking Residual Network 50 (ResNet50) [12]. Unlike the conventional approach, which relies on tuning hyperparameters to research global optima, ResNet50 is beneficial in optimization. It uses skip connection that could smooth the loss landscape. The model could avoid local minima, and easily reach to the global optima [22, 25]. However, ResNet50 has a higher complexity that is likely to cause a high variance problem. We therefore scale down ResNet50 by decreasing the number and width of convolutional layers. The resulting model has narrower and shorter network architecture than ResNet50 and is named ResNet29 (Figure 2).

**Figure 2.**
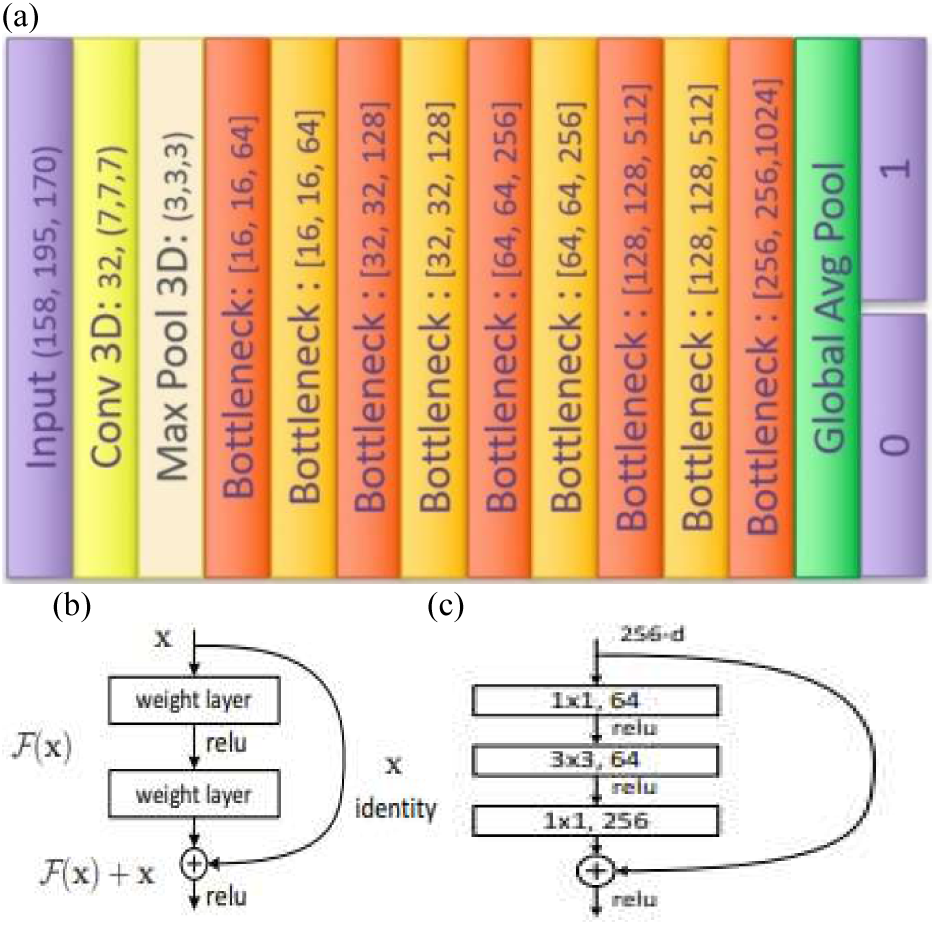
(a). Architecture of Convolutional Neural Network (CNN). The original ImageNet Model, i.e., ResNet50 was scaled down by narrowing and shortening the model. A Global Average Pooling layer was added at the end of the architecture, followed by the classification layer. (b) Skip connection was used to enable the model to reach a global optima. [12] (c). Bottleneck layers were set to reduce the model’s complexity and thereby improve the classification performance. [12]

ResNet29 is an end-to-end binary classification model. The number of filters in the first convolutional is reduced from 64 to 32. The number of bottleneck modules in each convolutional section is reduced from 3, 4, 6, and 3 to 2, 2, 2, and 2, respectively. One additional bottleneck layer is added at the end. The number of filters of each residual block is divided by 4, resulting in 4,305,666 hyperparameters.

All codes are built in python Keras as TensorFlow backend. Experiments are conducted by using 4 NVIDIA P100 Pascal (12G HBM2 memory). The source task is completed in approximately 9 hours and 3 hours is taken to complete the target task.

### E. Hyperparameters

At the source task, the model is trained with a cyclically changing learning rate from 1e-2 to 1e-4 with a unit epoch of 25 through the entire epoch of 75 to avoid the model being stuck in local optima, in order to achieve global maxima [30]. To reduce overfitting, ridge regression and weight constraint with the hyper-parameter value of 4e-4 and 2 are used with the batch normalization layer [16]. To reduce gradient exploding, gradient clipping is set as 1 [26]. The model and the weight matrix trained on NC vs. DAT are transferred to the target task of classifying MCI-NC vs. MCI-C. At the target task, the first 127 out of 155 layers are frozen during training, which results in 2,767,106 trainable parameters. The model is retrained with a cyclically changing learning rate from 1e-3 to 1e-5 with unit epoch 25 through the entire epoch 125. Ridge regression, weight constraint, and gradient clipping are set as 7e-4, 2, and 1 with batch normalization layer.

Batch size is fixed at 1. All convolutional layers are initialized with ‘*he_normal’* [13] and the ‘*elu’* activation function is used while output layer uses ‘*softmax’* activation function. Categorical cross entropy is used as a loss function and stochastic gradient descent is used as an optimizer.

### F. Feature Visualization Method: Occlusion Map

An occlusion map is generated by the prediction score of the model. For all subjects that have been corrected predicted by the target task, their preprocessed brain scan are occluded by a 2×2×2 voxel patch (intensity 0) then fed into the model. The patch position is iterated through each voxel with stride of 2. An occlusion map of prediction is generated and visualized as a heatmap. In places where prediction score decreases from the un-occluded result, these regions are regarded as significant in their contribution to the prediction of conversion to DAT.

### G. Relating to Clinical and Neuropsychological Measures

For relating 3D-CNN prediction probability to diagnostic and clinical measures, we include the Clinical Dementia Rating-Sum of Boxes (CDRSB), Alzheimer’s Disease Assessment Scale – cognitive 11 item (ADAS 11) and cognitive 13 item (ADAS 13), Mini Mental State Exam (MMSE), Rey Auditory Verbal Learning Test (RAVLT) – RAVLT Immediate, RAVLT Learning, RAVLT Forgetting, RAVLT Percent Forgetting, and Functional Activities Questionnaire (FAQ) [11, 23, 24, 27, 28, 36].

For 514 MCI-NC and 277 MCI-C subjects (Figure 1.), we calculate Pearson correlation coefficients between model prediction probabilities from the first MCI-diagnosed sMRI scan and rate of change in clinical assessments. The longitudinal clinical scores from the first MCI-diagnosed time point to the end of clinical history are used to obtain the rate of change of clinical assessments scores. Correlations between the baseline sMRI scan and the clinical scores’ rate of change obtained through the first to the last clinical history are also examined.

## V. RESULTS

### A. 3D-CNN classification results

Classifying MCI-NC vs. MCI-C through transfer learning with a base model of ResNet29 is successful (See Table 2.). It produces a test set classification accuracy of 82.4% and 0.827 Area Under the Curve (AUC) as well as 0.189 Equal Error Rate (EER) value (Figure 3.)

**Table 2.** Summary of MCI-NC vs. MC-C classification research.

| Table 2. Summary of MCI-NC vs. MC-C classification research. |  |  |  |  |  |  |
| --- | --- | --- | --- | --- | --- | --- |
|  | Engineering | Biomarker | Conversion time (years) | Random guess (%) | Accuracy (%) | Increase (%) |
| **Proposed model** | **Network** | **sMRI** | **3** | **50.7** | **82.4** | **31.7** |
| Basaia et al, 2018 | Feature | sMRI | 3 | 65.6 | 74.9 | 9.3 |
| Suk et al, 2017 | Feature | sMRI, Clinical Score | 1.5 | 57.5 | 74.8 | 17.3 |
| Cheng et al, 2015 | Feature | sMRI, PET, CSF | 2 | 56.6 | 79.4 | 22.8 |
| Li et al, 2014 | Feature | MRI, Meta features<sup>3</sup> | 4 | 54.9 | 74.8 | 19.9 |

**Figure 3.**
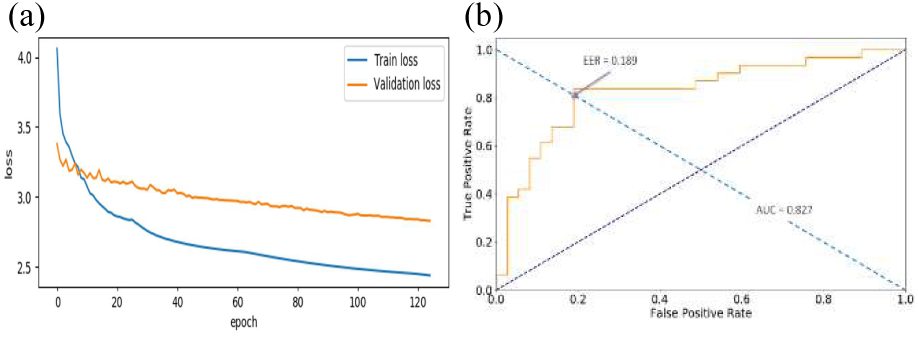
Loss history of train and validation data (a) and classification performance (b), i.e., Area Under the Curve (AUC) and Equal Error Rate (ERR) on test data. Train and validation loss are continuously decreasing along to the epochs, which indicates the model is learning. Weight matrix that is restored and used to evaluate the test classification accuracy was where the validation loss showed the minimum. Test classification accuracy reported 82.4%. AUC and EER value are 0.827 and 0.189, respectively.

The test set is composed of MCI-C patients whose conversion time is between 0 to 3 years. To further look at the models’ prediction performance over a longer conversion time, a separate MCI-C dataset whose conversion time is longer than 3 years is used. In conversion time from 0 to 3 years, 3 to 6 years, and 6 to 10 years, there are 37, 39, and 9 MCI-C subjects, and the model’s sensitivity on these three groups are 79.31, 70.27, and 55.57, respectively. The same model and its produced weight matrix are implemented to predict patients with longer conversion time. The results show that prediction score decreases with longer conversion time (Figure 4).

**Figure 4.**
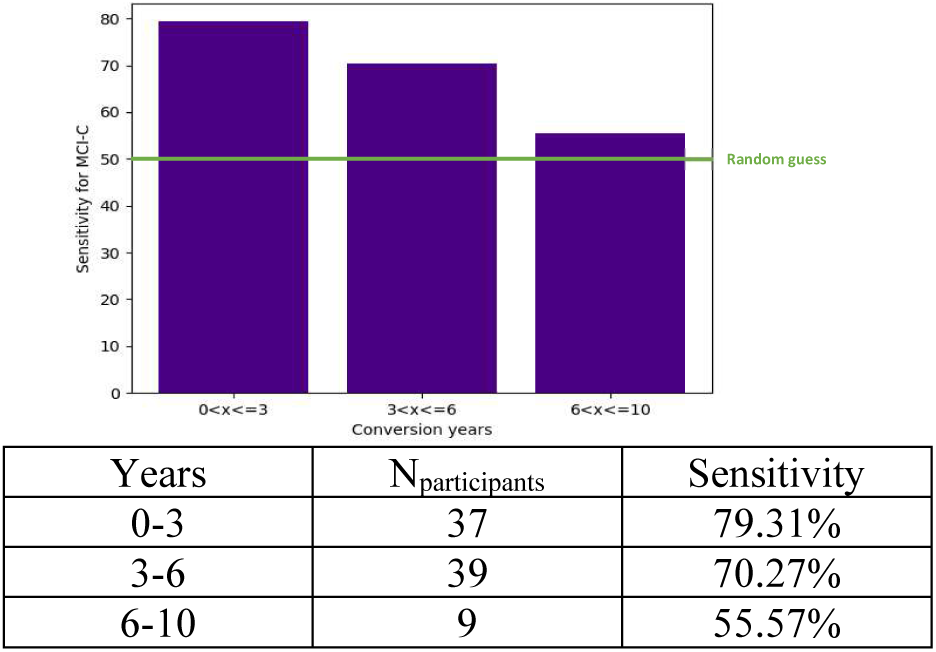
The sensitivity to predict patients with conversion years from 0 to 10. The same model and its weight matrix show decreasing sensitivity as conversion time gets longer. It indicates that the heterogeneous nature of DAT makes the model confused in predicting future development.

### B. Feature Visualization

Using occlusion mapping, we identify structural features predicted by the model (Figure 5.). Red regions in the brain indicate a high prediction score, indicating a greater likelihood of being the MCI-NC brain. Blue regions indicate that the occlusions of these regions lower the model’s confidence in predicting MCI-NC status; As seen in Figure 5, the hippocampus, amygdala, and pons regions are relevant for characterizing MCI-NC Similarly, the cerebellum and pons regions are recognized as features in predicting MCI-C. classification (Figure 6).

**Figure 5.**
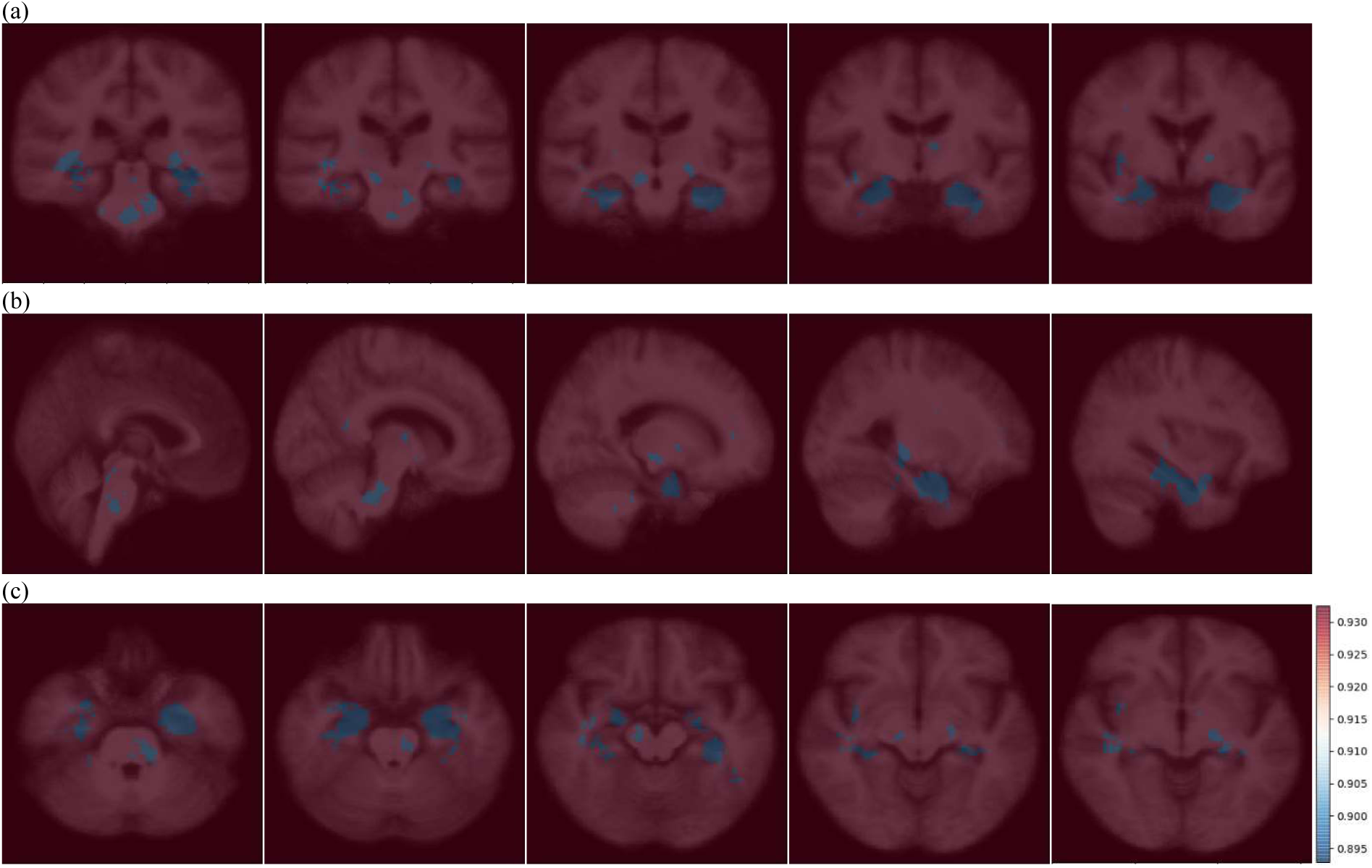
Occlusion map’s (a) coronal plane, (b) sagittal plane, and (c) transverse plane across all correctly predicted MCI-NC patients. The red color indicated the higher prediction score, whereas the blue color indicated the lower prediction score. The blue regions, which implies the important brain regions in predicting MCI-NC, indicate the hippocampus, amygdala, pons regions, etc.

**Figure 6.**
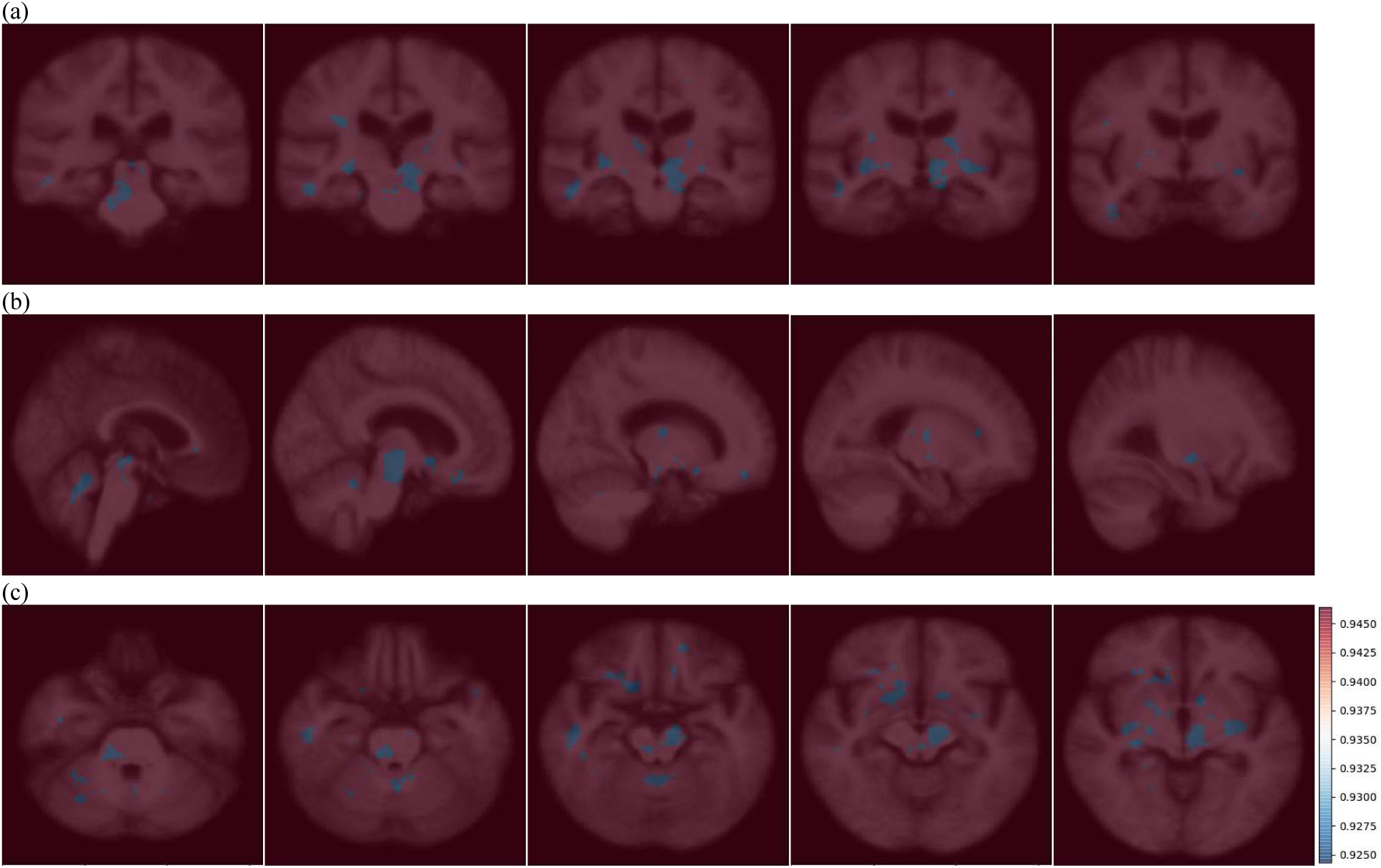
Occlusion map’s (a) coronal plane, (b) sagittal plane, and (c) transverse plane across all correctly predicted MCI-C patients. The red color indicated the higher prediction score, whereas the blue color indicated the lower prediction score. The blue regions, which implies the important brain regions in predicting MCI-C, indicate the cerebellum and pons regions, etc.

### C. Relating to *clinical scores*

CNN-based prediction score shows significant correlation with CDRSB, FAQ, MMSE, and RAVLT forgetting (Figures 7. and 8.). Higher prediction score of CNN is related to the higher score of CDRSB and FAQ and lower MMSE and RAVLT forgetting score. On the other hand, RAVLT immediate learning, ADAS11, and ADAS 13 do not show a significant correlation with the 3D-CNN-based prediction scores.

**Figure 7.**
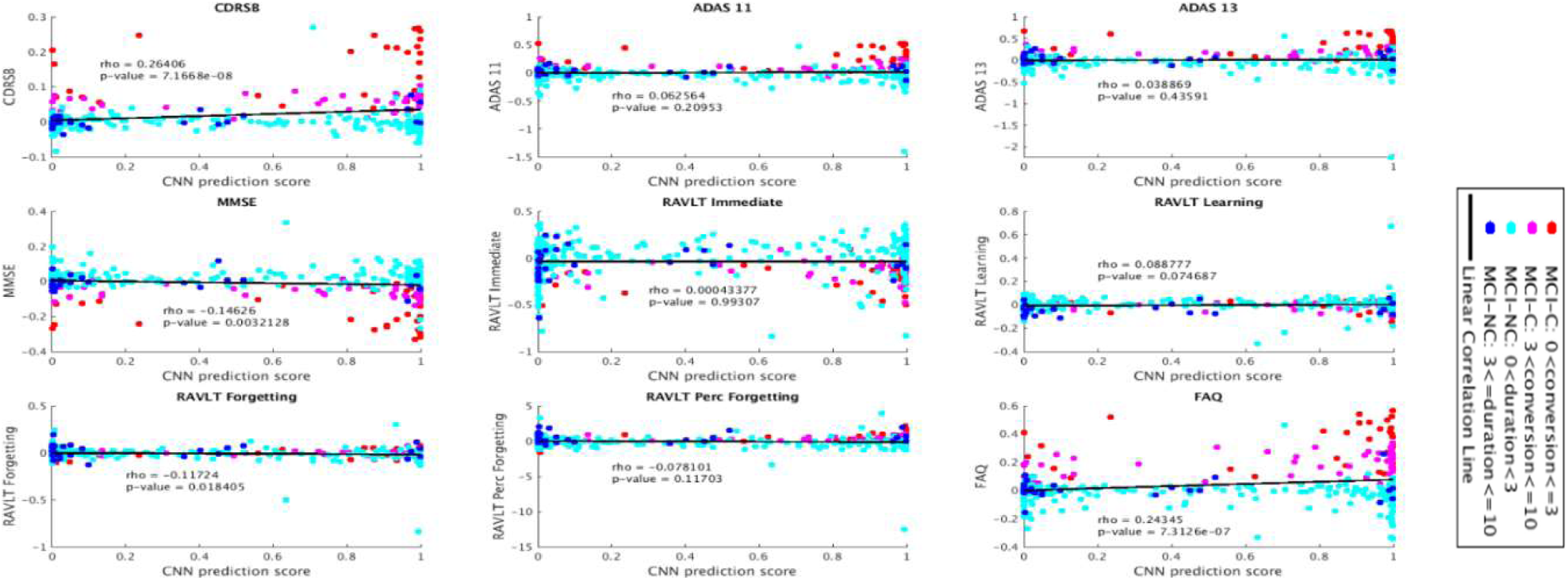
Correlation between CNN-based score from first MCI-diagnosed sMRI scan and clinical assessment scores’ rate of change.

**Figure 8.**
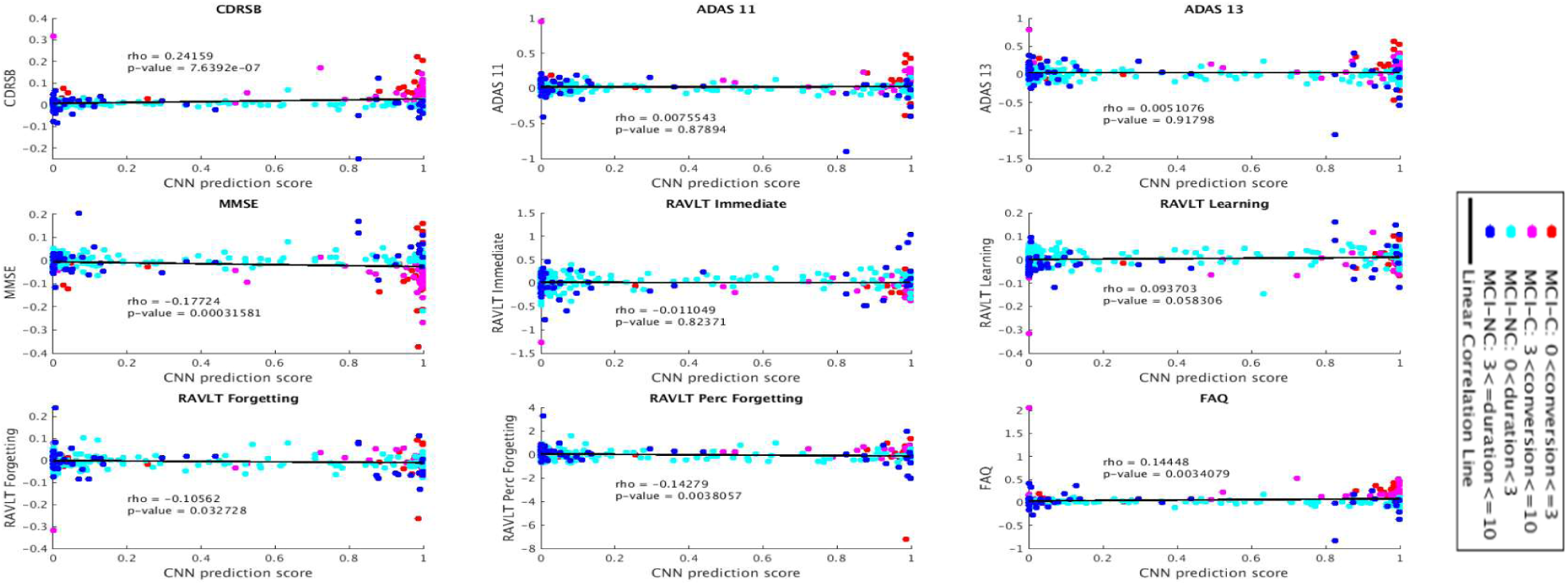
Correlation between CNN-based score from baseline sMRI scan and clinical assessment scores’ rate of change.

## VI. DISCUSSION

Using an end-to-end 3D-CNN deep learning model with transfer learning on structural MRI scans, we are able to predict MCI patients that either remained stable in their diagnosis (MCI-NC) or progressed to DAT (MCI-C) with an 82.4% accuracy. We achieve this without feature engineering. Furthermore, we utilize an occlusion map and show the hippocampus, amygdala, and pons are key regions in characterizing the MCI-NC and cerebellum and pons in characterizing MCI-C. Finally, we show that prediction scores from our model are related to worsening of clinical and neuropsychological performance measures.

One of the latest experiments predicting MCI-NC vs. MCI-C defined the conversion time at 3 years [4], and we chose this conversion time for the present study in order to directly compare performance. Further, setting conversion time at 3 years provides a well-balanced data set between MCI-NC (N=222) and MCI-C (N=228) [15].

Compared to previous studies (Table 2), our model achieves the highest accuracy (82.4%) in classifying MCI-NC from MCI-C, and the largest difference from chance. It also shows the most significant prediction increase from random guess, i.e., 31.7%. It should be noted that conversion times in previous studies range from 1.5 years [38], 3 [4], to 4 years [21], while the present study uses a 3-year conversion time window.

Some existing studies also include multimodal biomarkers in their prediction models, such as positron emission tomography (PET) and cerebrospinal fluid (CSF) data [7]. Our model outperforms these models, demonstrating that it is possible to predict conversion to DAT by using a single sMRI scan. This improved performance of our model is due to the source task in the transfer learning scheme that provides as much diverse domain knowledge as possible. Also, various engineering techniques, for example, cyclically changing learning rate [30], and carefully-tuned a set of hyperparameters contributes to the improvement of the classification performance.

The ability for deep-learning models to identify anatomical brain regions in predicting conversion from MCI to DAT, to the best of our knowledge, has not been demonstrated previously. The occlusion map identifies regions including the hippocampus, cerebellum, amygdala, and pons as significant. We note that the patch color (black) used in the occlusion map does not alter the visualization results, as we found identical results using a white colored patch. Interestingly, while volumetric changes of limbic structures in DAT are well documented in the literature [4, 15, 46], disease-related volumetric changes of brain stem structures (including pons) in patients with DAT are less well documented [47]. However, previous research has shown Braak-stage dependent changes in locus coeruleus, a noradrenergic nucleus located in the pons [48]. Neuropathological changes in the AD are associated with degeneration of the noradrenergic projections from the locus coeruleus, and cytopathology in this region has been highlighted as an early event predicting disease progression in DAT [49].

Current clinical diagnostic criteria cannot accurately identify clinical stages of MCI-NC and MCI-C [39]. Automated classification systems for MCI-NC vs. MCI-C, such as the method presented in this study, offer promise for informing the clinical prognosis of these patients. Furthermore, the methods presented here will be useful for identifying which patients would benefit most from selection into clinical trials. Our methods avoid problems faced in the field such as data shortage, high variance, and data leakage. Our research shows high accuracy in predicting conversion as well as novel visualization features, both critical to advancing our understanding of DAT.

## ACKNOWLEDGMENTS

This research was funded by grant AG055121 from the National Institute on Aging, and by grants from Brain Canada, CIHR, NSERC and Compute Canada.

ADNI data collection and sharing for this project was funded by the Alzheimer’s Disease Neuroimaging Initiative (ADNI) (National Institutes of Health Grant U01 AG024904) and DOD ADNI (Department of Defense award number W81XWH-12-2-0012). ADNI is funded by the National Institute on Aging, the National Institute of Biomedical Imaging and Bioengineering, and through generous contributions from the following: AbbVie, Alzheimer’s Association; Alzheimer’s Drug Discovery Foundation; Araclon Biotech; BioClinica, Inc.; Biogen; Bristol-Myers Squibb Company; CereSpir, Inc.; Cogstate; Eisai Inc.; Elan Pharmaceuticals, Inc.; Eli Lilly and Company; EuroImmun; F. Hoffmann-La Roche Ltd and its affiliated company Genentech, Inc.; Fujirebio; GE Healthcare; IXICO Ltd.; Janssen Alzheimer Immunotherapy Research & Development, LLC.; Johnson & Johnson Pharmaceutical Research & Development LLC.; Lumosity; Lundbeck; Merck & Co., Inc.; Meso Scale Diagnostics, LLC.; NeuroRx Research; Neurotrack Technologies; Novartis Pharmaceuticals Corporation; Pfizer Inc.; Piramal Imaging; Servier; Takeda Pharmaceutical Company; and Transition Therapeutics. The Canadian Institutes of Health Research is providing funds to support ADNI clinical sites in Canada. Private sector contributions are facilitated by the Foundation for the National Institutes of Health (www.fnih.org). The grantee organization is the Northern California Institute for Research and Education, and the study is coordinated by the Alzheimer’s Therapeutic Research Institute at the University of Southern California. ADNI data are disseminated by the Laboratory for Neuro Imaging at the University of Southern California.

## Footnotes

1 93 ROI for each sMRI and PET, and 3 features from CSF are used.

2 Clinical scores and genetic information include CDR, ADAS11, ADAS13, MMSE, RAVLT immediate, RAVLT learning, RAVLT forgetting, RAVLT percent forgetting, FAQ, APGN1, APGN2, APOE2, APOE3, and APOE4.

3 MRI features indicates average cortical thickness, standard deviation in cortical thickness, volumes of cortical parcellations, volumes of specific white matter parcellations, and the total surface area of the cortex. And Meta features includes demographic, genetic information, baseline cognitive scores, and lab tests. 305 MRI features and 52 Meta features are used.

